# Early loss of endogenous NAD^+^ following rotenone treatment leads to Sarm1 activation that is ameliorated by PARP inhibition

**DOI:** 10.1101/2021.07.30.454548

**Authors:** Ankita Sarkar, Malinki Sur, Sourav Dutta, Puja Dey, Piyali Mukherjee

## Abstract

Sarm1 is an evolutionary conserved innate immune adaptor protein that has emerged as a primary regulator of programmed axonal degeneration over the past decade. *In vitro* structural insights have revealed that although Sarm1 induces energy depletion by breaking down NAD^+^, it is also allosterically inhibited by NAD^+^. However, how NAD^+^ levels modulate the activation of intracellular Sarm1 has not been elucidated so far. This study focuses on understanding the events leading to Sarm1 activation in both neuronal and non-neuronal cells using the mitochondrial complex I inhibitor rotenone. Here we report the regulation of rotenone-induced cell death by loss of NAD^+^ that may act as a “biological trigger” of Sarm1 activation. Our study revealed that early loss of endogenous NAD^+^ levels arising due to PARP1 hyperactivation preceded Sarm1 induction following rotenone treatment. Interestingly, replenishing NAD^+^ levels by the PARP1 inhibitor, PJ34 restored mitochondrial homeostasis and prevented subsequent Sarm1 activation in rotenone treated cells. These cellular data were further validated in *Drosophila melanogaster* where a significant reduction in rotenone mediated loss of locomotor abilities and reduced dSarm expression was observed in the flies following PARP inhibition. Taken together, these observations not only uncovers a novel regulation of Sarm1 induction by endogenous NAD^+^ levels but also point towards an important understanding on how PARP inhibitors could be repurposed in the treatment of mitochondrial complex I deficiency disorders mediated by Sarm1.

## Introduction

Mitochondrial complex I deficiency has been reported in a number of diseases with mitochondrial disorders and has been strongly linked to patients with Parkinson’s disease (PD) [1, 2]. Rotenone, a lipophilic insecticide, has been shown to be a potent inhibitor of mitochondrial complex I [3, 4]. Several studies have indicated that rotenone mediated cell death is associated with mitochondrial depolarization, DNA damage and ROS formation that corroborates with similar observation in other mitochondrial complex I deficiency disorders [5, 6,7,8]. So, how this mitochondrial dysfunction induced by rotenone correlates with cell death? Our previous study has indicated that the TLR adaptor protein sterile alpha and TIR motif containing 1 (Sarm1) plays a pivotal role in this process as Sarm1 knockdown showed a significant reduction in rotenone induced cell death in SH-SY5Y cells [9]. Further, a study by *Hughes et al* has revealed that Sarm1 inhibition can prevent axonal degeneration following rotenone treatment [10]. Sarm1 is a NAD^+^-hydrolyzing enzyme that upon activation results in loss of nicotinamide adenine dinucleotide^+^ (NAD^+^) resulting in severe energy depletion within the axons with subsequent induction of axonal degeneration [11,12,13]. Although Sarm1 has emerged as an important regulator of programmed axonal destruction, how the expression and activation of endogenous Sarm1 is regulated within the cell is not well defined till date. Further, the non-neuronal regulation of this protein is understudied although this protein has been shown to be expressed in the liver, kidney and immune cells [14]. Since Sarm1 and rotenone induced cell death seems to be intricately linked, we envisioned that this may serve as an excellent cellular model to understand how this protein is endogenously regulated during mitochondrial complex I deficiency disorders and whether a single event or a cumulative phenomenon acted as a biological trigger of Sarm1 activation within the cells.

Although Sarm1 has been shown to mediate rotenone induced cell death, it is not known how NAD^+^ levels modulates this process. Recent reports indicated that NAD^+^ acts as a key determinant of mitochondrial health and homeostasis [15, 16] and decides cell fate by participating in cell death networks like apoptosis, autophagy or parthanatos [17,18,19]. Under situations of stress that induces mitochondrial damage, a process termed ‘mitophagy’ is initiated to remove these damaged organelles and restore cellular homeostasis. It has been suggested that upon various cellular stresses accumulation of damaged mitochondria arising due to incomplete clearance by mitophagy is associated with rapid cell death in the absence of a functional autophagy machinery [20]. Studies on rotenone mediated autophagy are inconclusive [21] and it is unclear whether defective mitochondrial clearance by mitophagy plays a determining role in rotenone induced cell death. Further, how NAD^+^ status affects the overall bioenergetic balance within the cells that may drive key cellular processes like autophagy and apoptosis is not clearly defined.

The two main classes of enzymes that are responsible for maintaining intracellular NAD^+^ pool are the NAD^+^ consuming enzymes Sirtuins and PARPs [22,23,24]. While the short-term activation of PARPs triggering a DNA damage response is considered to be crucial for maintaining cellular homeostasis, prolonged activation of PARPs leads to rapid degradation of NAD^+^ and eventually cell death [25, 26]. Interestingly, rotenone induced oxidative DNA damage has been reported in a few studies with Parkinson’s disease (PD) patient samples [27, 28] that could lead to PARP1 hyperactivation and loss of NAD^+^ following rotenone treatment.

We report here that rotenone induced temporal regulation of NAD^+^ levels lead to a sequential triggering of events that ultimately cause cell death. Our study indicates that rotenone treatment results in an early loss of NAD^+^ that is accompanied by mitochondrial dysfunction and PARP1 hyperactivation that may serve as the “biological trigger” of Sarm1 induction which has been previously shown to be allosterically inhibited by NAD^+^ using *in vitro* systems [29,30,31]. In concurrence with these observations, replenishing NAD^+^ levels by pre-incubation of cells with the PARP inhibitor PJ34 or Olaparib restored cellular NAD^+^ levels, prevented Sarm1 activation and significantly reduced cell death following rotenone treatment. To obtain a generalized overview of rotenone-mediated NAD^+^ regulation and Sarm1 activation, we conducted our studies in both non-neuronal and neuronal cell lines HEK293 and SH-SY5Y, respectively. To strengthen our hypothesis, we extended our findings in a previously established rotenone model of Drosophila and our results showed reversal of rotenone-induced locomotor deficits in the presence of the PARP inhibitor Olaparib in the flies. Thus, results generated from this study not only provides the first evidence of regulation of Sarm1 expression by endogenous NAD^+^ levels but also offers the exciting opportunity of repurposing PARP inhibitors in the treatment of Sarm1-induced pathological conditions and mitochondrial complex I deficiency disorders.

## Results

### Rotenone induced cell death in both non-neuronal and neuronal cells that was independent of caspase-3 activation

Mitochondrial complex I is the first point of entry of electrons to the electron transport chain and its inhibition has been implicated in different pathological conditions like PD [32]. Here we attempted to gain a comprehensive mechanistic insight in rotenone-mediated cell death in the routinely used non neuronal cell line HEK293 and the neuronal cell line SH-SY5Y. Our results indicate that rotenone induced cell death in a dose and time-dependent manner in both HEK293 and SH-SY5Y cells (Fig. 1 A and B). We observed that in HEK293 cells almost 40% cell death was achieved at a dose as low as 500 nM in 24 h. In comparison to the HEK293 cells, a dose of 5 μM could induce a comparable cell death phenomenon in the neuronal cell line SH-SY5Y, at 24 h (Fig. 1B) that also led to retraction of the cellular processes at 24 h post treatment (Fig. 1C). Due to its greater susceptibility to mitochondrial complex I inhibition, we further evaluated the status of apoptosis in HEK293 cells following rotenone treatment. Rotenone treatment in HEK293 cells resulted in caspase-3 activation as indicated by increased accumulation of active caspase-3 in the cells treated with 500 nM of rotenone (Fig. 1D). However, contrary to our expectation, prior incubation with the PAN caspase inhibitor Z-VAD-FMK followed by rotenone treatment did not reverse rotenone-induced cell death in both HEK293 and SH-SY5Y cells (Fig. 1E) indicating that caspase activation was perhaps not the sole factor driving rotenone induced cell death. This prompted us to perform an in-depth analysis of the underlying mechanism leading to rotenone induced cell death using both non-neuronal HEK293 cells and the neuronal SH-SY5Y cells.

**Fig. 1.**
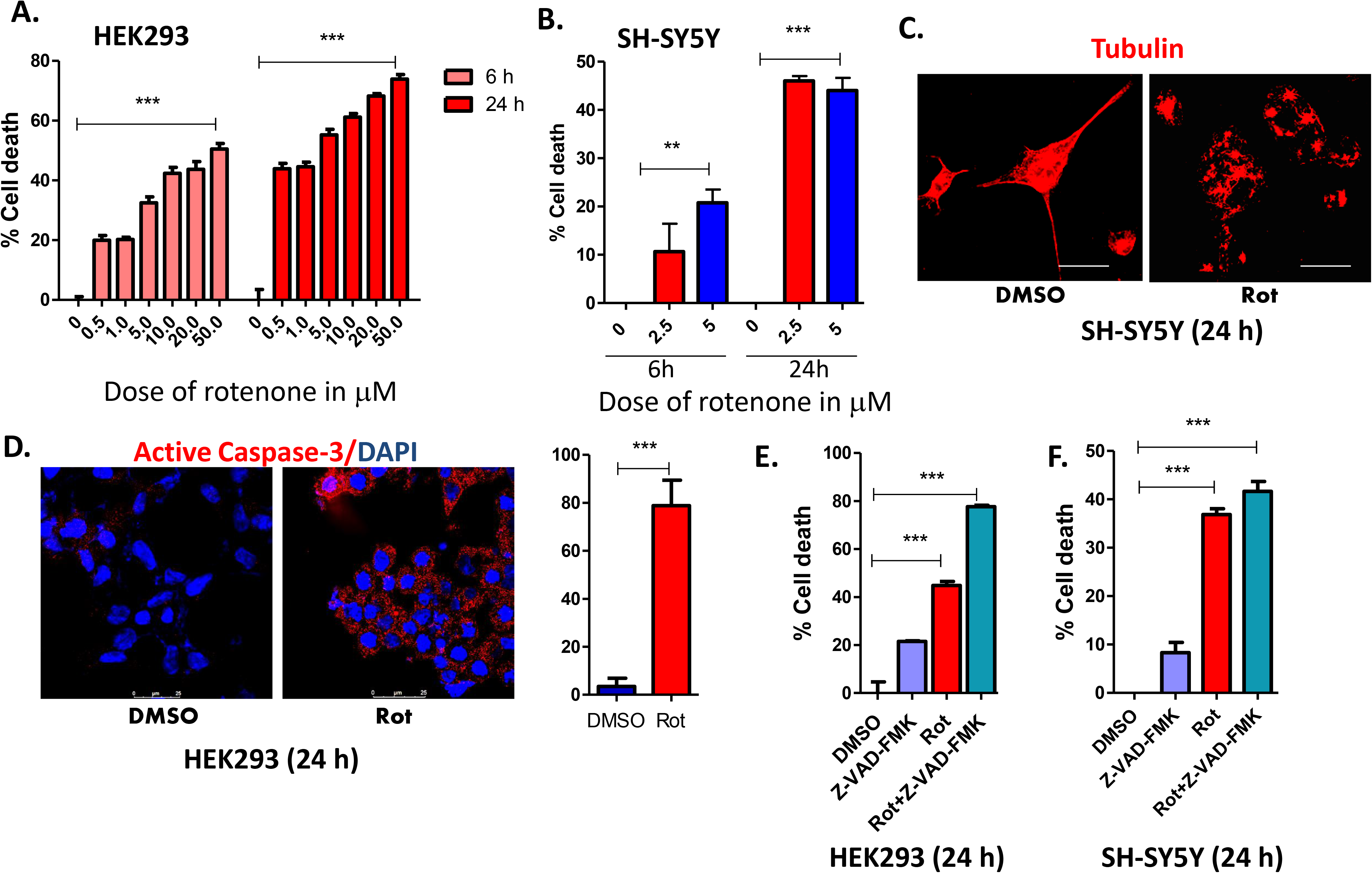
Rotenone induced cell death in both non-neuronal and neuronal cells that was independent of caspase-3 activation. **(A and B)** MTT assay of cells treated with indicated doses of rotenone in HEK293 (A) and SH-SY5Y (B) cells at 6h and 24h post treatment **(C)** SH-SY5Y cells treated with 5 µM of rotenone for 24 h and stained with anti-tubulin antibody to visualize the neurite processes following rotenone treatment. The results were compared with DMSO control samples. Scale bar represents 10 μm **(D)** Representative confocal images of HEK293 cells stained with anti-active Caspase-3 antibody (red) and DAPI (blue) in the presence of rotenone or DMSO control. Scale bar represent 25 μm. The right panel represents the percentage of active-caspase-3 positive cells in the presence of DMSO or 500 nM rotenone and results are representative of at least five individual fields per treatment **(E and F)** HEK293 (E) and SH-SY5Y cells (F) were pre-incubated with the pan-caspase inhibitor Z-VAD-(OMe)-FMK (20 µM) for 1 h prior to rotenone (500 nM or 5 μM respectively) treatment and cell viability was measured by MTT assay. All results are representative of atleast three independent experiments.

### Rotenone induced cell death is accompanied by early mitochondrial dysfunction, increased ROS accumulation and defective autophagy

Cytochrome c released from damaged or leaky mitochondria stimulates the cleavage of caspase-3 to its active form [33]. Since we observed caspase-3 activation we undertook a detailed analysis of mitochondrial status following rotenone treatment. Our results in HEK293 cells show mitochondrial membrane depolarization in these cells as early as 4 h post-rotenone treatment that preceded cell death as observed by staining with the cell permeant dye, TMRM that accumulates only in active mitochondria (Fig. 2A). A similar phenomenon was also observed in SH-SY5Y cells at 4 h post-treatment with 5 μM of rotenone (Fig. 2B). Further, live staining with mitotracker green (that localizes to the mitochondria irrespective of mitochondrial membrane potential) showed significant loss in mitochondrial puncta structure (as indicated by arrows in the enlarged panel) in rotenone treated SH-SY5Y cells (Fig. 2C) as well as in HEK293 cells (Fig. 2D) but the mitochondria stained brighter compared to untreated controls which could be indicative of accumulation of damaged mitochondria in these cells. It is well established that mitochondrial damage leads to increased ROS production [7] and accumulating ROS level causes further mitochondrial damage that may perpetuate ROS production in rotenone treated cells. Rotenone has been shown to induce cellular ROS generation but the source of this ROS is not well defined in HEK293 cells. To understand the source of ROS in these cells, we analyzed the status of mitochondrial ROS and observed an increase in mitochondrial ROS levels as indicated by increased Mitosox staining (Fig. 2E) as early as 4 h post-treatment. Conforming to previous reports, we observed an increase in total ROS levels in rotenone treated HEK293 (Fig. 2F) in a time-dependent manner and prior incubation with the anti-oxidant NAC significantly (*P* < 0.001) reduced ROS generation both at 6 and 24 h post treatment (Fig. 2F) in these cells. Interestingly, although there was heightened ROS production following rotenone treatment, prior incubation with NAC did not significantly (P value 0.1) reverse the ongoing cell death process in both HEK293 cells (Fig. 2G) as well as SH-SY5Y cells (Fig. 2H). Taken together, these data indicated that heightened ROS production may be the effect of accumulation of damaged mitochondria and may trigger a sequence of events that cannot be abrogated by prior treatment with NAC. It is interesting to note here that although mitochondrial dysfunction was observed as early as 4 h post rotenone treatment, apoptosis was not induced at this time point as no AIF accumulation was observed within the nucleus at 4 h post-treatment (Fig. 2I)

**Fig. 2.**
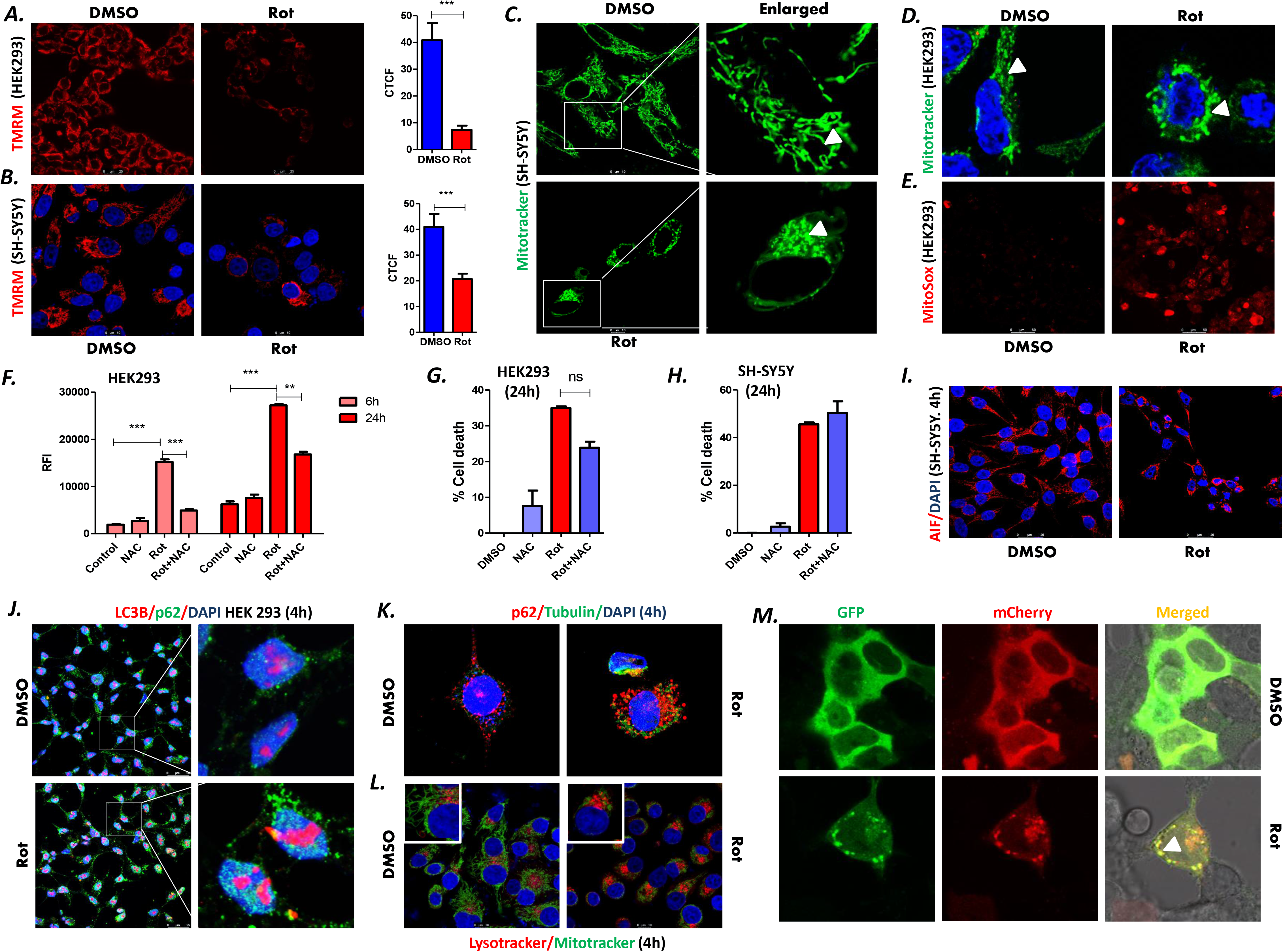
Rotenone induced cell death was accompanied by mitochondrial dysfunction, increased ROS accumulation and defective autophagy. **(A)** (Left panel) Immunofluorescence analysis of HEK293 cells incubated with 100 nM TMRM to assess the mitochondrial membrane depolarization in DMSO control and rotenone treated cells at 4 h post-treatment. Scale bar represents 25 μm. (Right panel) Corrected total cell fluorescence (CTCF) values of TMRM intensity of control vs rotenone treated cells **(B)** Immunofluorescence analysis of SH-SY5Y cells stained with 100 nM of TMRM at 4 h post rotenone treatment and compared with DMSO control. Scale bar represents 10 μm. (Right panel) CTCF values of TMRM intensity of control vs rotenone treated cells **(C and D)** Immunofluorescence analysis of SH-SY5Y (C) and HEK293 (D) cells stained with 100 nM of Mitotracker Green at 4 h post rotenone treatment and compared with DMSO controls. Scale bar represents 10 μm (C) and 5 μm (D). Arrows indicate intact or disrupted mitochondrial punctas in the enlarged panel (C) or in D **(E)** Visualization of mitochondrial ROS production in HEK293 cells by staining with the mitochondria specific ROS indicator Mitosox red at 4 h post rotenone treatment. Scale bar represents 50 μm **(F)** Relative fluorescence intensity (RFI) as a measure of ROS production in HEK293 cells with and without pre-incubation with 1mM NAC followed by rotenone treatment. Total ROS level was measured by H2DCFDA assay and fluorescence intensities were measured at Ex/Em: 492/527 nm **(G and H)** MTT assay of HEK293 (G) and SH-SY5Y (H) cells in the presence or absence of NAC pre-treatment followed by rotenone treatment for 24 h **(I)** Immunofluorescence analysis of SH-SY5Y cells treated with rotenone for 24 h and stained with anti-AIF (red) antibody and DAPI (blue) to visualize the nuclear localization of AIF post rotenone treatment and compared to DMSO controls. Scale bar represents 25 μm. **(J)** Immunofluorescence assay of HEK293 cells stained with anti-LC3B (red), anti-p62 (green) antibodies and DAPI (blue) to compare the expression and localization of the autophagy proteins followed by rotenone treatment. Scale bar represents 25 μm in the left panel. The indicated cells are cropped to show the single cell localization of these proteins **(K)** Immunofluorescence assay of SH-SY5Y cells with anti-p62 (red), anti-Tubulin (green) antibodies and DAPI (blue) to assess p62 accumulation following rotenone treatment (5 μM) for 24 h **(L)** Immunofluorescence analysis of SH-SY5Y cells co-stained with mitotracker green and lysotracker red at 4 h post rotenone treatment to compare the colocalization of the mitochondria with the lysosome. Scale bar represents 10 μm (upper panel) and 6 μm (lower panel). **(M)** Autophagic flux determination in HEK293 cells with transient transfection with mCherry-EGFP-LC3B plasmid for 24 h followed by rotenone treatment for 4 h

Accumulation of damaged mitochondria that resulted in increased ROS production could be due to a defective mitochondrial turnover via autophagy. Therefore, to correlate mitochondrial dysfunction and increased ROS production in HEK293 cells, we were inclined to check the status of autophagy in these cells. Microtubule-associated protein 1 light chain 3B (LC3B) and p62/sequestosome 1 are two widely used autophagy markers [34, 35]. Our results showed an increase in p62 accumulation (an indicator of defective autophagic flux) in rotenone treated HEK293 cells as well as SH-SY5Y cells as early as 4 h post-treatment (Fig. 2J and K). Co-staining with endogenous LC3B showed higher nuclear LC3B staining in rotenone treated HEK293 cells as compared to the control cells indicated by arrows (Fig. 2J right panel). Further, co-staining with mitotracker green and lysotracker red did not reveal any co-localization in rotenone treated cells at 4 h post-treatment (Fig. 2L) which indicated that mitophagy was not initiated in these cells at this earlier time point. We next sought to understand whether p62 accumulation following rotenone treatment was due to decreased autophagosome-lysosome fusion or defect in autophagic flux. For this, cells were transfected with the mCherry-GFP-LC3 plasmid and since GFP signal is lost under the acidic environment of the lysosome, the mCherry positive cells confirms a functional autophagic flux within the cells [35, 36]. Transfection with mCherry-GFP-LC3 showed increased accumulation of yellow puncta in rotenone treated cells as opposed to the red puncta (Fig. 2M) indicating it is not the autophagosome formation but a defect in autophagic flux that could result in limited turnover of damaged mitochondria in rotenone treated cells. Real time analysis of key autophagy genes revealed several fold higher inductions of the Atg genes (Fig. S1A-C) as compared to the DMSO treated control cells but reduced expression of a few late autophagy genes like Ulk1 (unc-51-like kinase 1) that plays an important role in the formation of autophagosome at both early and late time points following rotenone treatment in HEK293 (Fig. S1A) and SH-SY5Y cells (Fig. S2B-C).

### Rotenone treatment resulted in depletion of cellular NAD^+^ levels and PARP inhibition by PJ34 prevented rotenone-induced cell death in both HEK293 and SH-SY5Y cells

Since rotenone-induced autophagy did not go to completion, we asked whether energy deficits within the cells prevented autophagic degradation of damaged mitochondria via the lysosomal pathway. Accumulating recent evidence indicates that cellular NAD^+^ homeostasis is intricately associated with functional autophagy [37, 38] which helps the cells further to adapt to energy deprivation by obtaining energy from autophagic degradation products. Here we speculated whether the NAD^+^-consuming enzymes of the cells like PARP1 and Sirt1 have any role in autophagic defect that may lead to increased accumulation of damaged mitochondria and drive rotenone mediated cell death. Our results suggested that rotenone treatment reduced NAD^+^/NADH ratio thus decreasing total cellular NAD^+^ levels in HEK293 cells at 24 h post rotenone treatment (Fig. 3A). Interestingly, prior incubation of cells with the PARP1 inhibitor PJ34 significantly (P<0.001) reversed rotenone induced cell death in HEK293 cells (Fig. 3B). To further confirm the importance of NAD^+^ replenishment in the reversal of rotenone induced cell death, we analyzed the role of PJ34 in the neuronal cell line SH-SY5Y and a significant (P<0.001) restoration of cell death following prior incubation with PJ34 was also observed in these cells (Fig. 3C). However, a dose dependent analysis revealed that a higher does (50–100 μM) of PJ34 was required to achieve this reversal as compared to 25 μM in HEK293 cells. There was also a differential expression pattern of the genes involved in NAD^+^ metabolism pathway especially that of *Nmnat1* and *Nmnat2* (key enzymes implicated in the regulation of the NADase Sarm1 mediated axonal degeneration) in rotenone treated HEK293 cells (Fig. S2A-D). Notably, loss of NAD^+^ was observed as early as 2 h in rotenone treated HEK293 cells (Fig. 3) and pre-incubation of these cells with PJ34 followed by rotenone treatment restored cellular NAD^+^ levels at 4 h post treatment indicating that early drop in NAD^+^ levels was a key player in rotenone induced cell death. In comparison to the PARP inhibitor, the Sirtuin1 inhibitor EX527 had no such effect on rotenone mediated cell death (Fig. 3E) in HEK293 cells. To understand whether PJ34 had a similar effect on other inhibitors of the mitochondrial OXPHOS system, HEK293 cells were treated with the mitochondrial complex III and V inhibitor Antimycin and Oligomycin respectively which revealed that PJ34 had no effect in the reversal of cell death induced by these inhibitors (Fig. S2 E-F) and was specific to the complex I inhibitor rotenone.

**Fig. 3.**
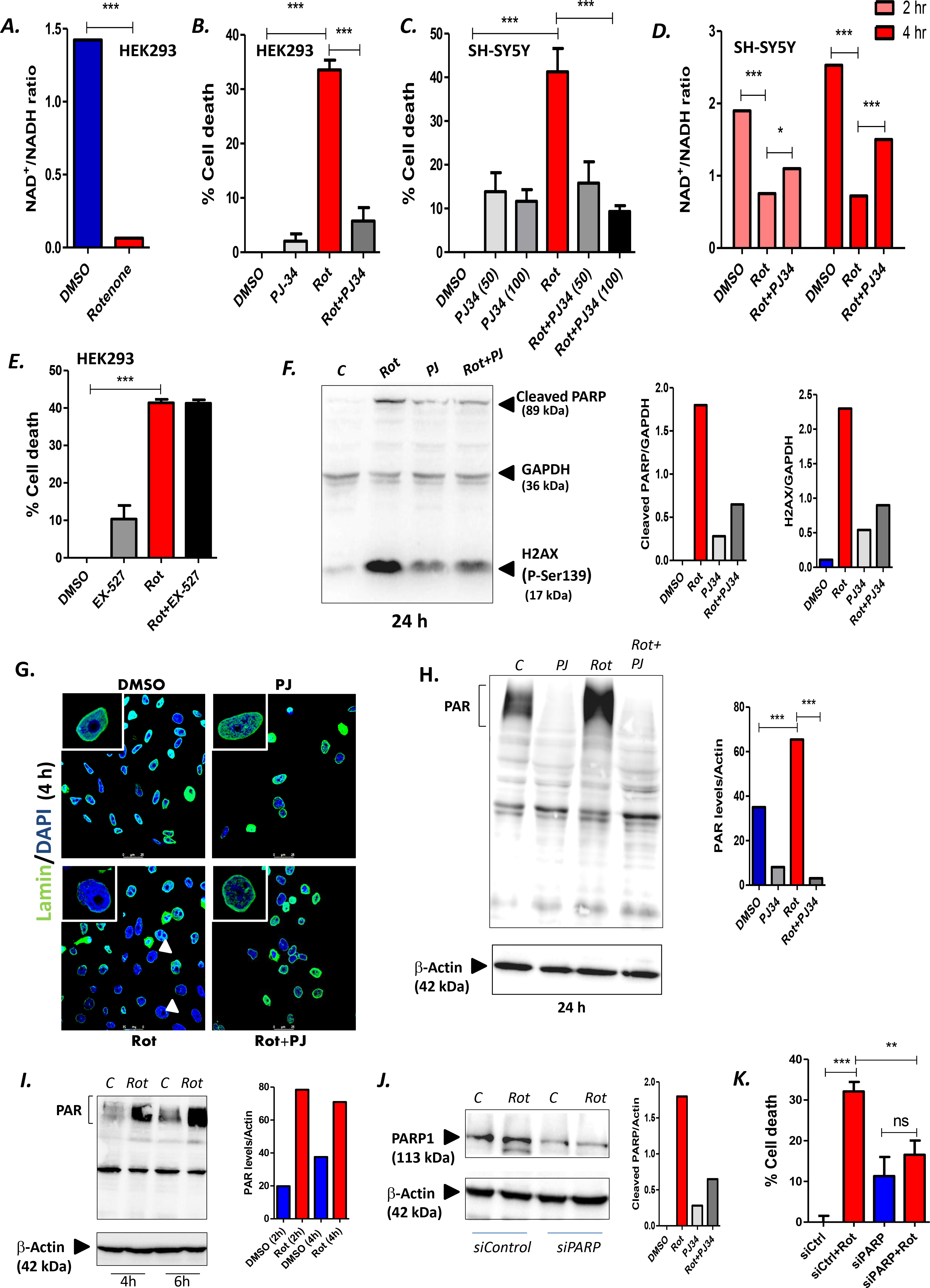
Rotenone treatment resulted in depletion of cellular NAD^+^ levels and PARP inhibition by PJ34 prevented rotenone-induced cell death in both HEK293 and SH-SY5Y cells. **(A)** Determination of NAD^+^/NADH ratios by fluorometric assay in HEK293 cells 24 h post rotenone (500 nM) treatment and compared with DMSO treated controls **(B)** Cell viability was measured using MTT assay in HEK293 cells pre-incubated for 1 h with 25 µM of the PARP inhibitor PJ34 followed by 500 nM of rotenone treatment for 24 h and the results were compared with DMSO control **(C)** Cell viability was measured using MTT assay in SH-SY5Y cells pre-incubated for 1 h with 50 or 100 µM of the PARP inhibitor PJ34 followed by 5 µM of rotenone treatment for 24 h and the results were compared with DMSO control **(D)** NAD^+^/NADH ratios by fluorometric assay in SH-SY5Y in an early time point to show the rapid loss of NAD^+^ post rotenone treatment and results were compared with samples pre-incubated with PJ34 (50 μM) to determine the role of PARP inhibition **(E)** Cell viability assay in the presence of Sirt1 inhibitor EX-527 (25µM) in HEK293 cells. Cells were pre-incubated with EX-527 for 1 h followed by rotenone treatment for 24 h **(F)** (Left panel) Immunoblot analysis of DNA damage and apoptosis marker proteins H2AX and cleaved Parp1 respectively in rotenone treated cells with or without prior treatment with PJ-34. Gapdh served as loading control. A cocktail antibody containing the proteins mentioned were used (Right panel) Densitometric analysis to determine the relative protein levels with respect to the loading control was performed in Image J software **(G)** Representative confocal images of SH-SY5Y cells pre-incubated with PJ34 (50 μM) and immuno-stained with anti-Lamin(A/C) antibody (green) and DAPI (blue) to visualize the nuclear lamin disintegration at 4 h post rotenone treatment and the results were compared to DMSO and PJ34 only controls **(H)** (Left panel) Immunoblot analysis of PARP1 hyperactivation by determining the PARylation levels. (Right panel) Densitometric analysis of PAR (Poly ADP Ribose chain) to determine the relative level of PAR in indicative treatments **(I)** Immunoblot analysis of PARP1 hyperactivation in SH-SY5Y cells at the indicated time points. Densitometric analysis of PAR (right panel) were performed in the samples to determine the relative level of PAR **(J)** (Left panel) Immunoblot analysis of PARP1 to determine the level of knock-down by siRNA in SH-SY5Y cells (Right panel) Densitometric analysis of siRNA knock-down samples for PARP1. β-Actin serve as loading control for immunoblot analysis and the quantification performed by Image J software. **(K)** Cell viability assay in PARP1 knock-down SH-SY5Y cells followed by rotenone treatment for 24 h and the results were compared with control siRNA samples. Data are representative of at least three independent experiments per panel.

PARP is a DNA damage sensor and it has recently been shown that there is increased localization of PARP1 to H2AX.2 enriched chromatin damage sites [39] and phosphorylated H2AX.2 is a histone marker of DNA double strand breaks [40]. Our results showed that there was increased level of phosphorylated H2AX (p-Ser 139) in the presence of rotenone, the level of which is significantly decreased following pre-incubation with PJ34 and subsequent rotenone treatment (Fig. 3F). This reduction in DNA damage alongwith the reversal mediated by PJ34 suggested that reduction of PARP hyperactivity in the presence of rotenone may prevent early NAD^+^ loss thus reducing cellular energy deficits. Additionally, the nuclear lamins which plays an integral role in the maintenance of genome integrity and induction of apoptosis was also disrupted in the presence of rotenone, a phenomenon that was reversed in the presence of PJ34 (Fig. 3G).

To confirm DNA-damage induced PARP hyperactivation, we next sought to analyze the level of poly (ADP-ribose) (PAR) formation in rotenone treated cells [41]. As indicated in Fig. 3H, there was a significant elevation of PAR formation at 24 h post-rotenone treatment that was dramatically prevented in the presence of PJ34. Next, we performed a time-dependent analysis of PAR formation in rotenone treated cells to determine whether PARP hyperactivation preceded apoptosis and correlated with the early drop in NAD^+^ levels. A robust increase in PARylation was observed as early as 4 h following rotenone treatment (Fig. 3I) that strongly suggested that ROS emanating from damaged mitochondria could result in sustained DNA damage and PARP hyperactivation thus reducing NAD^+^ levels in rotenone treated cells that is prevented in the presence of PJ34. To further understand the importance of the role of PARPs in rotenone-induced cell death, we conducted a siRNA mediated knockdown of PARP1 in SH-SY5Y cells (Fig. 3J) and analyzed cell death in the presence of rotenone (Fig. 3K). There was a significant (P value 0.0024) reversal of cell death following knock down of siPARP in rotenone treated cells (Fig. 3K) further pointing towards the importance of PARP1 inhibition to prevent rapid NAD^+^ loss in rotenone mediated cell death.

### Prior incubation of cells with PJ34 prevented the activation of the NADase Sarm1 following rotenone treatment

Previous reports suggested that the NADase Sarm1 is required for rotenone induced cell death [9, 10] but the mechanism remains poorly understood. Here we observed increased Sarm1 expression at 24 h post-rotenone treatment in the HEK293 cells but there was no induction of Sarm1 at the earlier time points in both HEK293 and SH-SY5Y cells (Fig. 4A and B) indicating that mitochondrial dysfunction and deregulation of autophagy machinery occurred prior to Sarm1 induction in these cells. We also observed increased Sarm1 expression in 3T3 cells (Fig. S3A) but the levels were several folds lower than HEK293 cells which may account for greater resistance of these cells to rotenone induced cell death (Fig. S3B). Interestingly, transfection of HEK293 cells with siSarm1 resulted in partial reversal of rotenone induced cell death (Fig. S3C) that indicated that Sarm1 could be a primary mediator of rotenone induced cell death in non-neuronal cells as well. Although one of the Sarm1 antibodies was previously standardized in western blot assay in primary neurons, these antibodies yielded non-specific bands in western blot assay of HEK293 cells. Thus, in order to confirm Sarm1 knockdown by siRNA transfection, we performed a real time analysis of the Sarm1 gene which showed a significant reduction in Sarm1 expression (Fig. S3D). We further evaluated Sarm1 protein levels in SH-SY5Y cells using an antibody that worked well in immunofluorescence studies in these cells and observed an increase in the Sarm1 protein levels at 24 h post rotenone treatment (Fig. 4C Lower Panel) compared to 4 h (Fig. 4C Upper Panel) further strengthening the observation that Sarm1 induction is a late event following rotenone treatment.

**Fig. 4.**
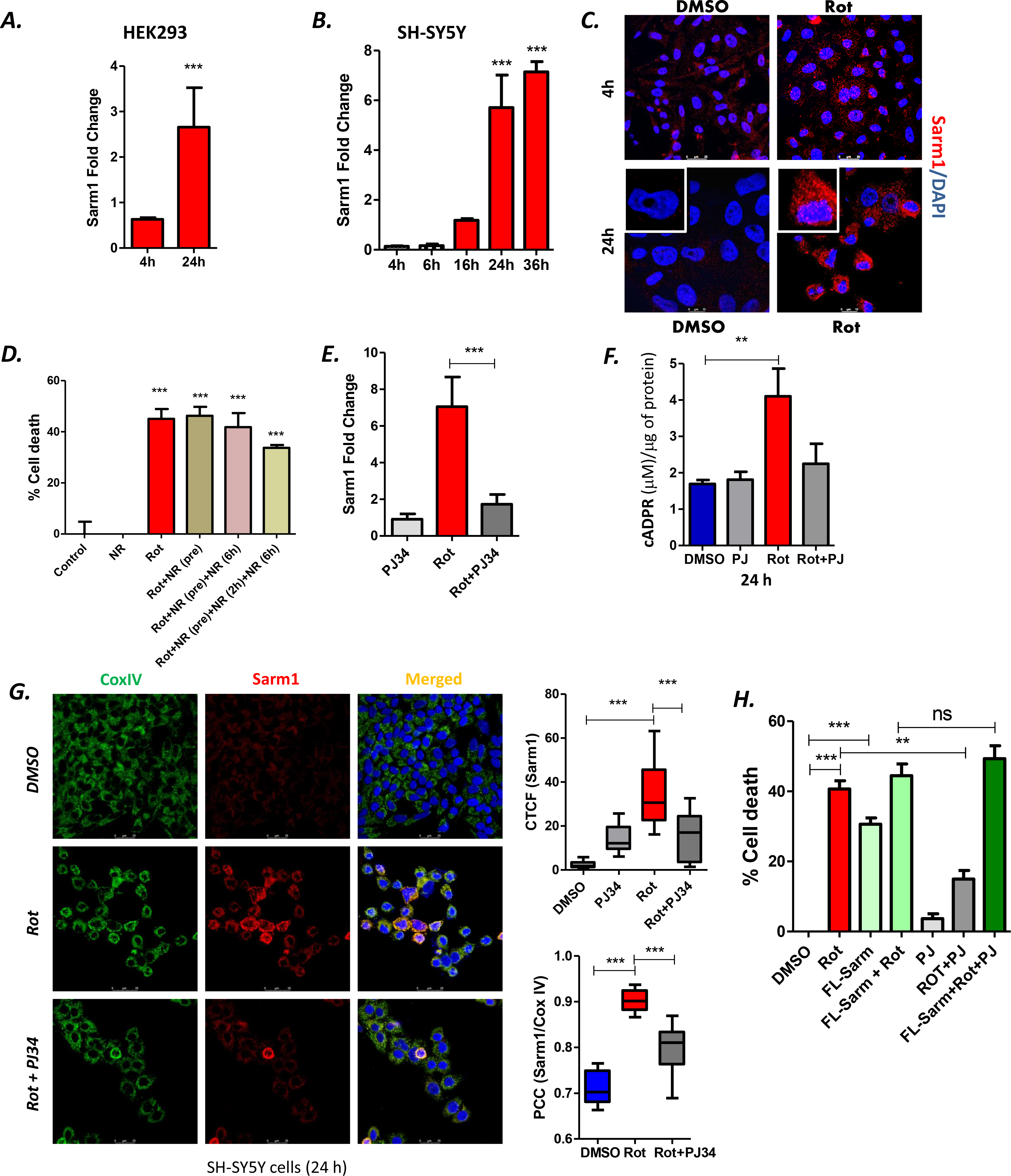
Prior incubation of cells with PJ34 prevented the activation of the NADase Sarm1 following rotenone treatment. **(A and B)** Relative fold change of mRNA expression of Sarm1 gene at 4 and 24 h post rotenone treatment in HEK293 (A) and at 4, 6, 16, 24 and 36 h post rotenone treatment in SH-SY5Y cells (B). Gapdh was used as an internal control **(C)** Immunofluorescence analysis of endogenous Sarm1 (red) in the presence or absence of rotenone at 4 h (Upper panel) and 24 h (Lower panel) post-rotenone treatment. Scale bar represents 25 μm for upper panel and 10 μm for lower panel. Nuclear staining is represented with DAPI (blue) **(D)** HEK293 cells were pre-incubated with 1mM NR followed by rotenone treatment for 24 h or intermittent replenishment with NR at the times indicated and MTT assay was conducted to analyze percent cell death. Data are representative of at least three independent experiments per panel **(E)** Relative fold change of mRNA expression of Sarm1 gene in rotenone treated (24 h) HEK293 cells by qRT-PCR with respect to DMSO treated control samples in the presence or absence of PJ34. Gapdh was used as an internal control **(F)** SH-SY5Y cells were treated with indicated treatments for 24 h and cADPR levels were quantified by cADPR assay kit and the concentration of cADPR was expressed as µM/μg of protein. Data are representative of at least three independent experiments per panel **(G)** Representative confocal images (left panel) of SH-SY5Y cells treated with rotenone in the absence or presence of PJ34 for 24 h and immunostained with anti-COXIV (green), anti-Sarm (red) and DAPI (blue) to visualize the localization and levels of Sarm1 at the mitochondria after indicated treatments. Comparative analysis of CTCF values was performed to quantify the intensities in each sample. CTCF values were determined for at least fifteen fields for each sample (upper right panel). The bottom right panel represents the colocalization of Sarm1 with COXIV. Pearsons’ coefficients were determined by Image J software for indicated treatments and compared with DMSO control **(H)** Cell viability assay in FL-Sarm1 overexpressed SH-SY5Y cells transfected for 48 h followed by rotenone treatment for 24 h in the presence or absence of PJ34 and the results were compared with control untransfected samples. Data are representative of at least three independent experiments per panel.

It has been previously shown that exogeneous addition of the NAD^+^ precursor nicotinamide riboside (NR) reverses axonal degeneration induced by overexpressed Sarm1 [42]. To understand whether NR had a similar effect on rotenone induced cell death in HEK293 cells, we pre-incubated the cells with 1 mM NR for 1 h followed by addition of rotenone for 24 h. Our results indicated that pre-incubation with NR had little or no effect on rotenone induced cell death (Fig. 4D). We hypothesized that NR is probably being rapidly used up and hence cannot combat the early NAD^+^ loss that is further exacerbated following Sarm1 activation. Hence, we exogenously added NR at 6 h to the cells that were previously pre-incubated with NR followed by rotenone treatment. We also supplemented NR at low doses (0.5 mM) at 3 h and 6 h to avoid the accumulation of NMN (nicotinamide mononucleotide) due to excess NR supplementation. However, supplementing NR at different time points and with different doses had no significant effect on rotenone induced cell death in the HEK293 cells (Fig. 4D). These results indicated the importance of early NAD^+^ loss induced by PARP1 hyperactivation in rotenone mediated cell death. This is consistent with the recent Cryo EM observations in vitro that loss of NAD^+^ may release the autoinhibition of the ARM domain of Sarm1 by NAD^+^ leading to triggering of its NADase activity [30]. In line with this observation, we next sought to analyze whether reversal of cell death and restoration of cellular NAD^+^ levels by PJ34 correlated with Sarm1 induction. Real-time PCR analysis of PJ34 treated cells followed by rotenone treatment resulted in a prominent decrease in expression of Sarm1 at 24 h post treatment (Fig. 4E). It has been demonstrated recently that cleavage of NAD^+^ by Sarm1 results in the production of cADPR [43, 44]. To understand whether the increased Sarm1 levels correlated with its activation, we measured the cADPR levels in the neuronal SH-SY5Y cells at 24 post rotenone treatment. We found a significant increase in cADPR levels following rotenone treatment that was reduced but not completely abrogated in the presence of PJ34 in these cells (Fig. 4F).

Since Sarm1 has been shown to stabilize on depolarized mitochondria [45], we proceeded to check its localization at the mitochondria using co-staining with the Cox IV antibody following rotenone treatment in the presence and absence of PJ34. We show here for the first time that rotenone treatment induced translocation of Sarm1 to the mitochondria at 24 h post treatment as compared to DMSO treated controls (Fig. 4G) (Right Upper Panel) which was confirmed by the Pearson’s correlation coefficient (PCC) where there was a significant increase in the mitochondrial co-localization of Sarm1 (Fig. 4G Right panel) in rotenone treated cells. In continuation to our previous observation, we further observed a reduction in the Sarm1 protein levels alongwith reduced mitochondrial localization in these cells in the presence of PJ34 (Fig. 4G). To check the status of Sarm1 protein levels in HEK293 cells, we transfected these cells with FLAG-tagged full length Sarm1 followed by rotenone treatment. Interestingly, rotenone treatment stabilized the Sarm1 protein in these cells at 24 h post treatment compared to the untreated DMSO control cells as evident from the more intense staining of the FLG-tagged Sarm1 (Fig. S3E).

To understand the importance of the temporal induction of Sarm1 as observed in both HEK293 and SH-SY5Y cells, we asked whether PJ34 could still be functional in preventing rotenone-induced cell death in cells overexpressing Sarm1. Our data suggests that in contrast to the un-transfected cells, PJ34 could not reverse rotenone induced cell death in Sarm1-overexpressed cells which point towards the importance of the early events leading up to Sarm1 activation that may act as the final trigger of cell death driving the cells towards a point of no-return.

### PJ34 restored mitochondrial homeostasis and prevented accumulation of damaged mitochondria in rotenone treated HEK293 and SH-SY5Y cells

Since we hypothesized that early mitochondrial damage could subsequently leads to PARP1 hyperactivation via ROS induced oxidative DNA damage leading to early NAD^+^ loss and Sarm1 induction, we checked the status of mitochondria in rotenone treated cells exposed to PJ34. Pre-incubation of cells with PJ34 prevented early loss of mitochondrial membrane potential (Fig. 5A) as early as 4 h post rotenone treatment. To further confirm the role of PJ34 in the maintenance of mitochondrial homeostasis, we co-stained rotenone treated HEK293 cells pre-incubated with PJ34 with Mitotracker green and Lysotracker red. Two populations of cells were observed at 4 h as indicated in Fig. 5B. Cells showed intact mitochondrial staining as indicated by mitotracker green staining in rotenone treated cells in the presence of PJ34 (Fig. 5B upper panel). However, it was interesting to note increased co-localization of mitotracker green and lysotracker red in these cells as indicated by arrows (Fig. 5B lower panel) that maybe due to improved lysosomal turnover of damaged mitochondria. This was also confirmed by the Pearson’s correlation coefficient (PCC) which showed higher mitochondrial co-localization with the lysosomes (Fig. 5B, right panel) following PJ34 treatment in the presence of rotenone. Restoration of mitochondrial puncta structure was also noted in PJ34 treated SH-SY5Y cells followed by rotenone treatment (Fig. 5C). We further observed a decrease in the mitochondrial ROS levels as early as 4 h post rotenone treatment in the presence of PJ34 (Fig. 5D) that was accompanied by a significant decrease in the total cellular ROS levels in these cells (Fig. 5E) that strengthened our hypothesis of increased ROS production via accumulation of damaged mitochondria induced by rotenone.

**Fig. 5.**
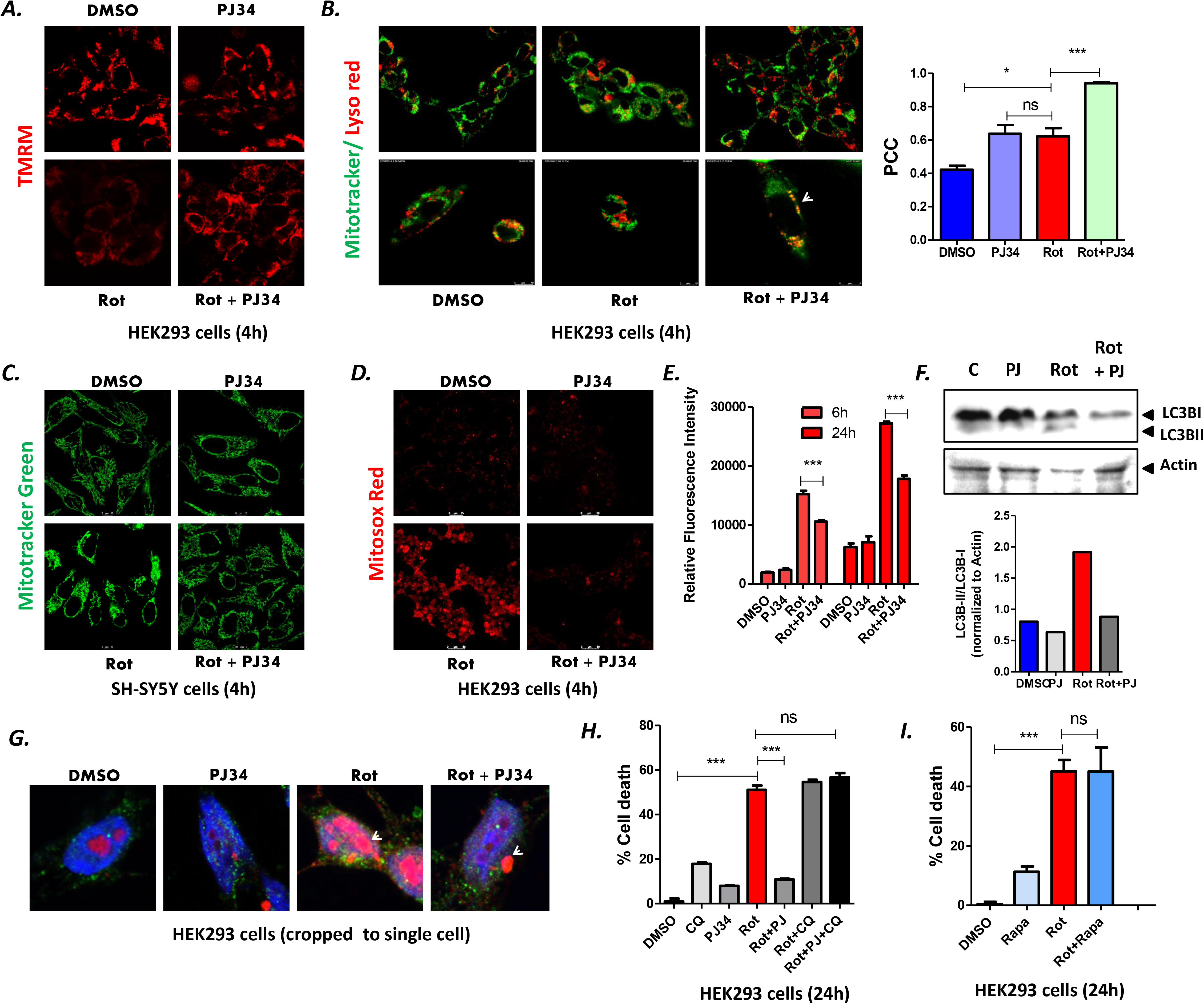
PJ34 restored mitochondrial homeostasis and prevented accumulation of damaged mitochondria in rotenone treated HEK293 and SH-SY5Y cells. **(A)** Immunofluorescence analysis of HEK293 cells incubated with 100 nM TMRM to assess the mitochondrial membrane depolarization in control and rotenone treated cells in the presence or absence of PJ34 at 4 h post rotenone treatment. Scale bar represents 25 μm **(B**) Immunofluorescence analysis of HEK293 cells co-stained with mitotracker green and lysotracker red at 24 h post rotenone treatment to compare the colocalization of the mitochondria with the lysosome in the presence or absence of PJ34. Scale bar represents 25 μm (upper panel) and 6 μm (lower panel). The arrow represents the colocalization in rotenone and PJ34 treated cells and the degree of co-localization was quantified by determining Pearson’s correlation coefficient (PCC) by Image J software (right panel) **(C)** Representative confocal images of SH-SY5Y cells incubated with mitotracker green (100 nM) to determine intactness of mitochondria with or without PJ34 pre-treatment followed by rotenone. Scale bar represents 25 μm **(D)** Visualization of mitochondrial ROS production in HEK293 cells by staining with the mitochondria specific ROS indicator Mitosox red at 4 h post rotenone treatment in the presence or absence of PJ34. Scale bar represents 50 μm **(E)** Measurement of total ROS levels by H2DCFDA assay and at Ex/Em: 492/527 nm by DCFDA assay to compare the overall ROS production in rotenone treated cells in the presence of absence of PJ34 **(F)** Immunoblot analysis of the autophagy marker protein LC3B in HEK293 cells treated with rotenone (24 h) in the presence or absence of PJ34. β-actin was used as a loading control **(G)** HEK293 cell co-stained with anti-LC3B (red) and anti-p62 (green) and nuclear stain DAPI (blue) to visualize the localization of autophagy marker proteins post rotenone treatment. Results were compared with DMSO control and PJ-34 treated cells. Single-cell cropped images are shown. **(H)** Cell viability assay in the presence of autophagic inhibitor chloroquine (50µM) with or without incubating the HEK293 cells with PJ34 prior to rotenone treatment **(I**) Cell viability assay in HEK293 cells pre-incubated with the **a**utophagy inducer Rapamycin (200 nM) prior to rotenone treatment for 24 h to determine the role of autophagy induction in response to rotenone treatment. Data are representative of at least three independent experiments per panel.

Since PARP inhibition by PJ34 improved mitochondrial turnover within the lysosomes which may subsequently lead to restoration of mitochondrial homeostasis [46], we sought to analyze the status of autophagy following PARP inhibition. Both Western blot and confocal microscopy analysis revealed a decrease in LC3B-II/I ratio (Fig. 5F) and reduced signal strength of LC3B (indicated by white arrows) in the nucleus (Fig. 5G) 4 h following rotenone treatment in the presence of PJ34. Since mitochondrial damage and turnover seems to be intricately linked, we sought to further understand whether functional autophagic machinery is required for PJ34 mediated protection from rotenone induced cell death. To test this hypothesis, cells were pre-incubated with chloroquine (an inhibitor of lysosomal acidification) and no reversal of cell death induced by rotenone was observed in the presence of a combination of chloroquine and PJ34 in HEK293 (Fig. 5H). Interestingly, stimulation of autophagy by rapamycin also could not overcome rotenone induced cell death (Fig. 5I). Collectively, these results suggest that a functional autophagy is intricately linked with cell death in rotenone treated cells that can be alleviated by PJ34 by restoring cellular NAD^+^ levels and cannot be achieved by stimulating autophagy alone.

### PARP inhibition by Olaparib prevented rotenone induced neurotoxicity and reversed rotenone induced locomotor deficits in 1-day old *Drosophila melanogaster*

In order to strengthen our understanding on the effect of PARP inhibition, apart from PJ34 treatment and siPARP1 mediated knockdown assays, we further extended our studies with another PARP inhibitor Olaparib. Although both PJ34 and Olaparib are known to inhibit PARP catalytic activity [47, 48], Olaparib has also been shown to induce PARP-DNA trapping [49] and induce DNA damage in a dose-dependent manner. Hence, before we proceeded to understand the effect of Olaparib in rotenone induced neurotoxicity, we first exposed SH-SY5Y cells to increasing dose of Olaparib and measured the induction of cell death. Our results revealed that a dose of 50 μM induced around 30% cell death in these cells. Hence, we chose a sub-optimal dose of 20 μM to study the effect of olaparib on rotenone induced cell death (Fig. 6A). Pre-incubation of cells with Olaparib followed by rotenone treatment significantly reversed rotenone induced cell death in SH-SY5Y cells at 24h post treatment (Fig. 6B). To further understand the physiological relevance of PARP inhibition by Olaparib in rotenone induced pathological conditions, we extended our data in the fruitfly *Drosophila melanogaster*. Olaparib has shown promise as an anti-cancer chemotherapeutic drug and is sold under the trade name Lynparza to treat ovarian cancer 33[50, 51]. Due to its known prospect in drug repurposing, we selected Olaparib for PARP inhibition instead of PJ34 in our fly studies.

**Fig. 6.**
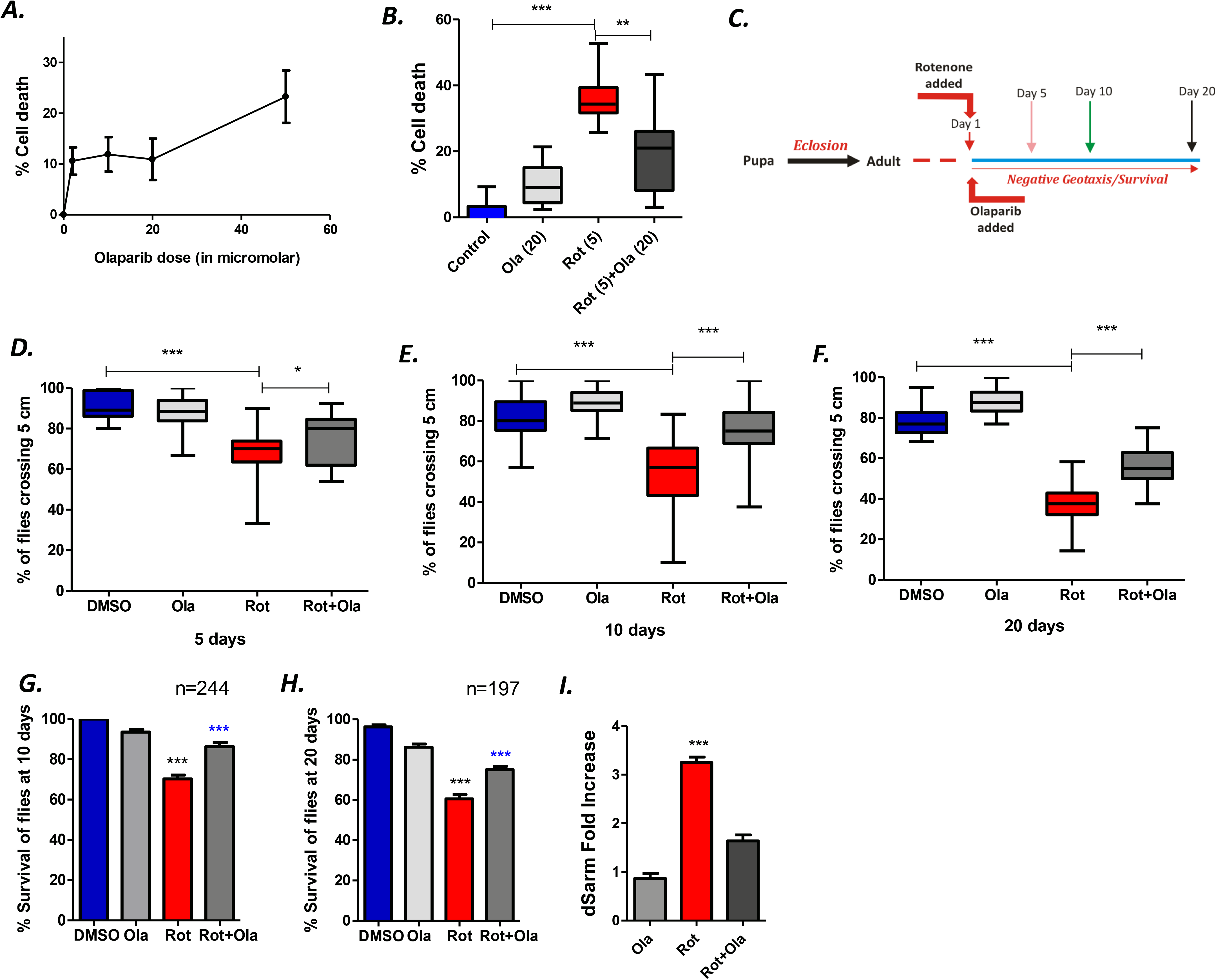
PARP inhibition by Olaparib prevented rotenone induced neurotoxicity and reversed rotenone induced locomotor deficits in 1-day old *Drosophila melanogaster.* **(A)** Dose dependent cell death analysis of Olaparib in SH-SY5Y cells at 24 h post-treatment. The percentage of cell death was determined by MTT assay to compare the cell death in indicated treatments **(B)** MTT assay of SH-SY5Y cells treated with rotenone (5 μM) in the presence or absence of 20 μM of Olaparib (pre-treated for 1 h prior to rotenone treatment) at 24 h post-treatment **(C)** Schematic representation of the experimental setup in *Drosophila melanogaster* (D-F) Negative geotaxis assay of 1-day old flies at 5 (D), 10 (E) and 20 (F) days post exposure with DMSO, olaparib (100 μM), rotenone (200 μM) and rotenone + olaparib **(G-H)** Survival assay of flies exposed as described in E-F. *** denotes significance between DMSO control and rotenone exposed flies and *** denotes significance between rotenone and rotenone + olaparib exposed flies. Results are representative of at least three independent set of experiments.

Our previous study has shown progressive locomotor deficits accompanied by dSarm induction in w^1118^ flies exposed to rotenone [9]. We first conducted a dose dependent analysis of Olaparib due to the unavailability of such data in flies and measured its effect on fly survival and selected the doses 50 and 100 μM for further analysis with rotenone. Our results indicated that 1-day old flies exposed to 100 μM Olaparib in the presence of 200 μM of rotenone (Fig. 6C) exhibited significant reversal in their climbing ability at 5, 10 and 20-days post exposure (Fig. 6D-F). We further observed heightened survival of flies in the presence of 100 μM Olaparib in the presence of rotenone at 10 and 20-days post exposure (Fig. G and H). To check the status of dSarm in these flies, we conducted a real time PCR analysis of the fly heads at 5 days post rotenone treatment in the presence or absence of Olaparib. Our results show that Olaparib prevented rotenone-mediated dSarm induction in younger flies. This is the first observation of the effect of PARP inhibition by Olaparib in the reversal of rotenone induced locomotor deficits in flies and further highlights the importance of PARP inhibitors in modulating rotenone induced pathological conditions.

## Discussion

NAD^+^ is a critical factor in the regulation of cellular homeostasis including DNA repair, mitochondrial function and autophagy [46, 52] but no single study has linked these processes together. Rotenone, a potent inhibitor of mitochondrial complex I activity, induces loss of mitochondrial membrane potential and stimulates the production of reactive oxygen species (ROS) but the time-dependent regulation of these processes has not been discussed. A few earlier studies have shown that addition of NAD^+^ reduces rotenone induced cell death [53] which is an important observation but how NAD^+^ regulate life/death decision following rotenone treatment is not known. These are important preliminary evidences linking cellular NAD^+^ levels with mitochondrial dysfunction ultimately leading to cell death especially in the context of mitochondrial complex I inhibition by rotenone. Recent studies on the NADase Sarm1 have implied that NAD^+^ breakdown by Sarm1 could be a key event in rotenone induced cell death. However, although the isoquinoline inhibitor of Sarm1 (DSRM-3716) could significantly reverse rotenone induced axonal degeneration, defects within the mitochondria persisted indicating that there could be other regulators of mitochondrial function in these cells which act in a concerted manner to stimulate cell death [10]. Taking cues from these observations, here we have attempted to dissect the temporal regulation of mitochondrial dysfunction, ROS production and Sarm1 induction and how NAD^+^ orchestrates these processes in rotenone induced cell death.

Mitochondrial health and homeostasis are vital towards cell survival and dysregulation of mitochondrial function has been linked to several diseases ranging from neurodegenerative disease to cancer [54] . Mitochondria are an important source of cellular ROS which mediates redox signaling at physiological levels [7]. However, accumulation of ROS leads to mitochondrial damage [20] which maybe perpetuated within the cells if these damaged organelles are not effectively cleared primarily via the process of autophagy (often termed as “mitophagy”). To understand the cause and effect of rotenone induced mitochondrial damage, we conducted a time-dependent analysis in HEK293 and SH-SY5Y cells and observed an early defect in mitochondrial homeostasis and heightened ROS production that could not be ameliorated by the ROS scavenger NAC. This clearly implied that accumulating mitochondrial damage overrode the scavenging capacity of NAC in these cells and the disintegrated but the intense staining pattern of the mitochondria further strengthened this observation in both HEK293 and SH-SY5Y cells. A further analysis clearly indicated that there exists a defect in autophagic flux following rotenone treatment that may lead to an ineffective clearance of damaged mitochondria. Apart from the early defect in autophagic flux, we observed an increased accumulation of endogenous LC3B within the nucleus of rotenone treated cells. Such localization was not observed in the overexpressed LC3 (mCherry-GFP-LC3) which formed puncta typical of autophagic vacuoles within the cytoplasm following rotenone treatment. Such enhanced nuclear localization of endogenous LC3 is previously unreported and is an interesting outcome of rotenone induced autophagic defect that needs to be explored further.

It has been recently highlighted that the oxidation state of the important cellular co-factor nicotinamide dinucleotide (NAD) undergoes alteration following metabolic stress [15] but how NAD^+^ levels may directly contribute to the driving of the autophagic flux or clearance of damaged mitochondria is not clearly defined. The only direct evidence linking NAD^+^ levels to mitochondrial homeostasis has been studied in the context of Werner’s syndrome where the authors demonstrate that augmentation of NAD^+^ leads to improved lifespan via the restoration of mitophagy [55]. Although we hypothesized that the NAD^+^ levels could drop following the activation of Sarm1 NADase activity, the timing of heightened Sarm1 expression and these early cellular defects could be not explained. Hence, we conducted a time dependent analysis of mitochondrial function and NAD^+^ status following rotenone treatment. Contrary to our expectations, we observed a loss of NAD^+^ levels as early as 2 h post rotenone treatment which did not match with the time of Sarm1 induction and this early loss of NAD^+^ could not be reversed by replenishing the NAD^+^ pool with prior incubation with nicotinamide riboside (NR), a precursor of NAD. NR has a short half-life and it could be rapidly used up in the cells. However, intermittent replenishment with NR only slightly reversed rotenone induced cell death strongly indicating that other factors contributed to the rapid NAD^+^ loss within these cells.

In order to understand what causes the early NAD^+^ loss in rotenone treated cells, we looked into other NAD^+^-consuming enzymes within the cell. Besides the NADase Sarm1 and CD38, the protein deacylase family of Sirtuins and PARPs are known NAD^+^ consuming enzymes. Hyperactivation of PARP1 has been shown to impede autophagy due to consumption of NAD^+^ and blocking Sirt1 function, a key regulator of autophagy [56]. Our results demonstrated PARP hyperactivation as early as 4 h post rotenone treatment that may account for rapid loss of NAD^+^ in rotenone treated cells. To delineate the function of this PARP hyperactivation, we asked whether PARP inhibitors could restore cellular NAD^+^ levels and prevent rotenone-induced cell death. Our results showed that prior incubation with the PARP inhibitor PJ34 but not the Sirt1 inhibitor EX527 significantly reversed rotenone induced cell death. Further, PJ34 not only prevented depletion of cellular NAD^+^ levels following rotenone treatment but also restored mitochondrial homeostasis and reduced ROS production. Importantly, abrogating this early accumulating energy depleted state prevented Sarm1 mRNA induction and a significant reduction in Sarm1 protein levels was observed in rotenone treated cells in the presence of PJ34. Sarm1 is known to possess an N-terminal mitochondrial localization signal and rotenone induced the translocation of Sarm1 to the mitochondria which is known to occur under stressed conditions that was significantly reversed in the presence of PJ34. Quite interestingly, recent Cryo EM studies on Sarm1 structure and function has shown the presence of an auto-inhibitory ARM domain that remains bound to NAD^+^ and prevents the activation of the NADase containing TIR domain of SARM1 [57]. The importance of PARP1 in rotenone induced cell death was also confirmed in siRNA studies in SH-SY5Y cells where there was significant reversal in cell death at 24 h. Thus, it is strongly possible that this early loss of NAD^+^ induced by PARP1 hyperactivation following rotenone treatment may overcome this auto-inhibition subsequently leading to the activation of Sarm1 and severe energy deficits within the cell.

We extended our studies to another PARP inhibitor, Olaparib which showed a similar reversal in cell death in SH-SH5Y cells. The physiological relevance of our observation in the cellular studies was strengthened by the fact that inhibiting PARP by Olaparib reversed rotenone induced locomotor deficits in *Drosophila melanogaster* that was accompanied by reduction in dSarm levels. Taken together, this is an important finding that correlates loss of endogeneous NAD^+^ levels with the expression/activation of intracellular Sarm1 in both neuronal and non-neuronal cell lines as well in flies. This study also provides a very important lead in the understanding the role of PARP inhibitors in modulating cellular energy homeostasis and mitochondrial function as depicted in the proposed model (Fig. 7). It is possible that early mitochondrial dysfunction induced by rotenone leads to ROS production and oxidative DNA damage that leads to PARP activation. Sustained mitochondrial damage in the absence of functional autophagy leads to ROS accumulation, PARP1 hyperactivation and further loss of NAD^+^. This loss of NAD^+^ may remove the allosteric inhibition of the ARM domain of Sarm1 and lead to activation of its TIR-NADase activity leading to further energy deficits and ultimately cell death.

**Fig. 7.**
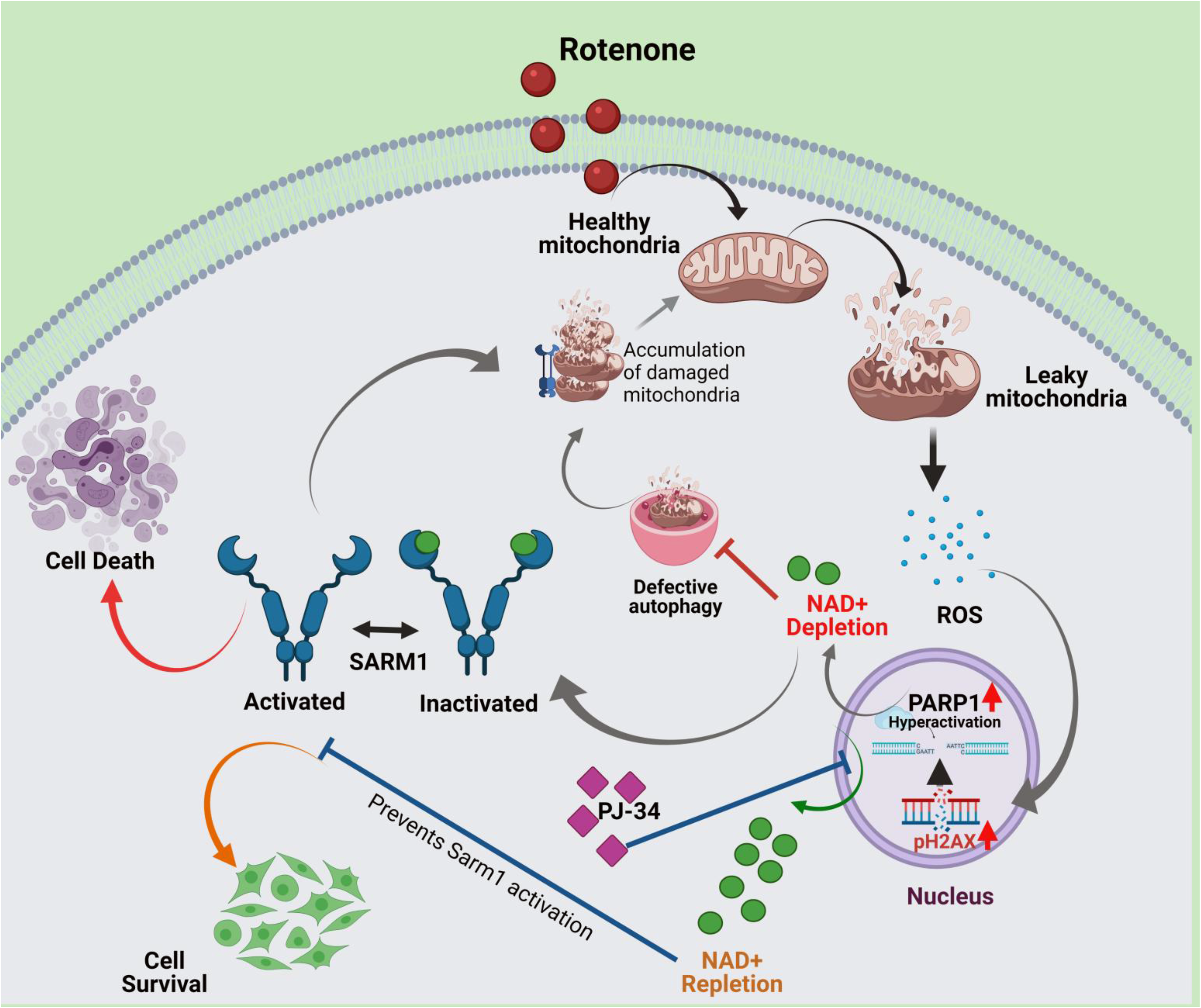
Proposed model of the study. Addition of rotenone to the cells results in mitochondrial depolarization and generation of reactive oxidative species (ROS). Excess ROS accumulation leads to nuclear DNA damage and PARP1 activation that leads to depletion of NAD^+^. This early loss of NAD^+^ blocks autophagic flux resulting in accumulation of damaged mitochondria, thus perpetuating further ROS production. In a vicious cycle this causes further DNA damage, PARP1 hyperactivation and subsequent NAD^+^ loss. This loss of NAD^+^ is predicted to remove the allosteric inhibition on the pro-apoptotic protein Sarm1 thus activating it. Sarm1 localizes to the depolarized mitochondria and induces subsequent cell death. Prior incubation with the PARP inhibitor, PJ34 replenishes cellular NAD^+^ levels, restores mitochondrial homeostasis and prevents Sarm1 upregulation in the cells thus preventing rotenone induced cell death. (*Created with BioRender.com*).

In conclusion, these observations could be an important step towards our understanding of the metabolic regulation of mitochondrial function, the autophagy-apoptosis network and Sarm1 activation by rapid loss of endogeneous NAD^+^ levels induced by the mitochondrial complex I inhibitor, rotenone.

## Materials and Methods

### Cell lines, antibodies and chemicals

HEK293 cells were obtained from the laboratory of Dr. Abhik Saha (Presidency University Kolkata, India), SHSY-5Y cells were obtained from Dr. Oishee Chakrabarti (Saha Institute of Nuclear Physics, Kolkata) and 3T3 cells obtained from Dr. Shubhra Majumder (Presidency University, Kolkata). The cells were maintained in Dulbecco’s modified Eagle’s medium (DMEM) (Gibco, Life Technologies) supplemented with 10% fetal bovine serum (FBS) (Invitrogen) and 1% Penicillin-Streptomycin Solution (Sigma).

Mouse monoclonal antibodies against β-actin (ab8224), p62 (ab56416) and LC3 A/B (ab128025), Cov IV (ab16056) and rabbit polyclonal antibody against anti-cleaved caspase-3 (ab2302) were obtained from Abcam. Rabbit polyclonal antibody for AIF (4642), PARP1 (46D11) were obtained from Cell Signaling Technology Inc., rabbit polyclonal antibodies against Sarm1 was obtained from GeneTex for western blot analysis and Sarm1 (D2M5I) for staining was obtained from Cell Signaling Technology Inc. Rabbit polyclonal antibodies against FLAG (F1804) was purchased from Sigma and for PAR (MCA-1480) was purchased from Bio-Rad. The antibody for the protein MANLAC2 (Lamin A/C) was developed by Morris, G.E. (DHSB Hybridoma product MANLAC2 (10F8)) and obtained from the Developmental Studies Hybridoma Bank, created by the NICHD and maintained at the University of lowa, Department of Biology, lowa City, IA 52242. In order to determine the DNA damage and apoptosis, the Apoptosis and DNA damage (H2A.X(S139) + cleaved PARP1 + Anti-GAPDH) Western Blot Cocktail (ab131385) was obtained from Abcam. Mouse and Rabbit polyclonal secondary antibodies were purchased from Abcam (ab97023 and ab97051 respectively).

Mitochondrial complex I inhibitor rotenone (Sigma), pan caspase inhibitor Z-VAD (OMe)-FMK (Abcam), PARP-1 inhibitor, PJ-34 hydrochloride (Abcam), SIRT1 inhibitor EX-527 (Abcam), autophagy inhibitor Chloroquine diphosphate salt (Sigma), the autophagy stimulator Rapamycin (Abcam) were used in this study at the indicated doses. Nicotinamide riboside (NR) was procured from ChromaDex, USA.

### Cell viability Assay

Cells were cultured in 24-well plates for 24 h and were treated with rotenone at the indicated doses and time. In some experiments, cells were pre-incubated for 1 h with either PJ-34, EX-527, Chloroquine, Bafilomycin, Rapamycin, Z-VAD-FMK or NR prior to the addition of rotenone. Wells in triplicate were incubated with the MTT reagent (Sigma) at the specific time points at a concentration of 5 mg/ml. Following 2 h of incubation, the MTT solution was aspirated and the cells were lysed in DMSO and the formation of formazan in each well was determined in a plate reader (BioTek) at an absorbance of 540 nm. All results were compared to DMSO treated controls or the solvent used for the assay.

### Plasmid and transfection assays

Cells were seeded at a density of 0.4x10^5^ cells/well, 24 h prior to transfection. Cells were transfected with pCMV6-Sarm1 with C-terminal Myc-DDK tag (a kind gift from Dr. Karin Peterson, Rocky Mountain Laboratories, NIAID, NIH, USA) or Flag-N-TIR-Sarm1 with Lipofectamine 2000 (Invitrogen), for 48 h using the manufacturer’s protocol. To measure the status of autophagy, HEK-293 cells were transfected with mCherry-GFP-LC3 (a kind gift from Rupak Datta, IISER Kolkata) for 24 hours followed by treatment with rotenone for 24 h.

siRNAs for PARP1 (#6304) were purchased from Cell Signaling Technology Inc. HEK293 and SH-SY5Y cells were seeded at a density of 0.4*10^6 in a 24-well plate, 24 h prior to transfection. HEK293 and SH-SY5Y cells were transfected with siRNA (100 nM) for 24 h followed by treatment with rotenone for 24 h. Cell viability was assessed in these cells as described previously and results compared with control siRNA (#6568) treated cells. Similarly, HEK293 and SH-SY5Y cells were transfected with 25 nM siSARM (s23032, Ambion, life technologies) for 48 h followed by 24 h rotenone treatment. All the siRNA silencing and plasmid transfection studies was done with lipofectamine 2000 purchased from Thermo Fisher Scientific.

### Whole cell extracts and Immunoblot Analysis

Cells were cultured in 6-well plates at a cell density of 0.8 x 10^6^ per well 24 h prior to treatment as indicated. Following treatment, cells were lysed in lysis buffer (150mM NaCl, 1% NP-40, 50mM Tris pH 8.0) containing a protease inhibitor cocktail (Pierce, Thermo Fisher Scientific) for the preparation of whole cell extracts. Protein concentration was estimated using the Bradford assay kit (Bio-Rad) and 15 μg of protein/well were resolved on 12% SDS-PAGE and subsequently analyzed by immunoblot analysis. Bands were detected by a chemiluminescent ECL system (Bio-Rad) and quantified by Image J software.

### Immunofluorescence assay

12 mm cover slips were placed in 24-well plates and coated with geltrex for 1 h before seeding. Cells were seeded at a density of 0.03 x 10^6^ cells/well 24 h prior to the treatment. Following treatment, cells were washed with 1x PBS and fixed with solution containing 4% paraformaldehyde and 0.2% Triton-X-100 for 10 min followed by blocking with 5% NDS buffer (normal donkey serum, 0.05% Triton-X-100 and 0.2 M Glycine) at room temperature for 30 min. The cells were incubated with primary antibody at 4°C overnight followed by incubation with secondary antibody (Alexa flour 488 or Alexa flour 594) for 1 h at room temperature. Nucleus was stained with DAPI (Bio-Rad) for 5 min along with secondary antibody and mounted with Fluoroshield Mounting Medium (Abcam) and sealed for analysis. In case of live staining the cells were seeded in grooved 35 mm plates (Thermo Fisher Scientific) at a density of 0.04 x 10^6^. After treatment, cells were incubated with Lysotracker red DND9 (Life technologies), Mitotracker green FM (Life technologies), TMRM (Sigma) and Mitosox (Life technologies) at 37°C for 15 min and then washed with 3x times in fresh DMEM followed by incubation with Hoechst 33342 (Sigma) for 5 min and further analyzed by confocal imaging. All images were acquired by DMi8 confocal laser scanning microscope (Leica-microsystems) and images were analyzed using image J software.

### NAD^+^/NADH Assay

The total NAD concentration, NAD^+^/NADH ratio and the concentration of total NAD^+^ and NADH was determined by NAD/NADH fluorometric assay kit from Abcam (ab176723) following manufacture’s protocol. The concentration of each component was calculated from the standard curve obtained by the standard solutions provided in kit. As per manufacturer’s instruction, respective concentrations were calculated by the equations described in the protocol that is Total (NAD/NADH) = (B/V) x D where B = total NAD/NADH amount in the sample well (µM), V=sample volume added in the sample wells, D= sample dilution factor. The NAD^+^ concentration was calculated with the same method from the same standard curve. NAD^+^/NADH assay results were obtained from three independent experiments for each sample.

### cADPR Assay

The concentration of cADPR was measured by Amplite Fluorometric cADPR-Ribose Assay Kit (20305) from AAT Bioquest. Briefly, SH-SY5Y cells were seeded at a density of 0.4x10^5^ cells/well and grown for 24 h prior to treatment. Cells were treated with 5 μM of rotenone for either 16 h or 24 h following which whole cell lysate was extracted as described previously. Total protein levels were estimated by Bradford reagent (Bio-Rad). Prior to the setting up the test experiment, a standard curve was prepared by the reagents provided in the kit. Standards, blank controls and samples were added to black well plates (50 µl) followed by the addition of 50 µl of ADPRC working solution in each well. After 1 h incubation at room temperature in the dark, NAD^+^ levels were detected by the addition of 40 µl of Quest fluor probe into each well followed by 40 µl of Assay solution II and incubation for 20 mins in dark followed by the addition of 30 µl of enhancer solution into each well. The fluorescence intensity was monitored at an excitation and emission of 420/480 nm. cADPR concentration for the test samples was expressed as µg of cADPR/μg of protein. cADPR assay results were collected from three independent experiment for each sample.

### Measurement of total cellular ROS

Intracellular ROS was measured by H2DCF-DA (Sigma, D6883) After treatment, cell lysate was prepared by NP-40 lysis buffer and H2DCFDA was added at a concentration of 50 µM and incubated for 1 h and fluorescence intensities were measured at Ex/Em: 492/527 nm.

### qReal-Time PCR

RNA samples were collected by standard Trizol (Sigma) method and cDNA was prepared by Bio-Rad iScript cDNA synthesis kit. The real-time data was analysed and quantified by CFX manager software (Bio-Rad). All the results were obtained from at least three independent experiments. The mRNA expression levels of every gene were quantified and normalized with reference gene. In case of some genes relative expression was plotted by ΔCT method to compare the basal level and level after treatment in every sample to show the difference in expression. In another case the expressions of genes expressed as fold change with respect to control samples by ΔΔCT method to determine the increase in the expression with respect to control samples. The RT-PCR results were analysed by Bio-Rad CFX manager. All the results were obtained from at least three independent experiments.

### Fly survival and negative geotaxis assay

W^1118^ flies were a generous gift from Dr. Rupak Dutta, IISER, Kolkata. Both male and female flies were grown at 22° C and 12-h light/dark cycle in incubator in standard corn meal-agar diet and changed every 3-5 days. Rotenone and Olaparib was freshly prepared as 100 mM stock solution in DMSO. Flies were transferred to rotenone or Olaparib containing fresh medium every 2-3 days to monitor survival and locomotor deficit.

For the survival assay 1-day old flies were transferred to DMSO, rotenone and rotenone-Olaparib containing fresh medium. The total number of flies alngwith the dead and viable ones were counted at 3-day interval and the percentage of survival was calculated up to 20 days for each set. Three different sets were prepared per experiment.

The negative geotaxis assay was performed in a similar manner to that published by Gargano et al (Gargano et al., 2005) with minor modifications. At least 20 treated and 20 control flies were transferred once a day in the early afternoon without anesthesia into a vertical glass climbing vial (length, 25cm; diameter, 3cm) to perform the assay. The climbing vials were placed in a rectangular frame in order to keep them upright. After a ten-minute acclimation period, the vials were tapped three times to initiate the negative geotaxis assay. After 10 s, the percentage of flies crossing the 5 cm mark was noted for each trial. This procedure was repeated a total of ten times. All behavioral experiments were performed at room temperature under standard light conditions.

### RNA isolation and qRT-PCR from fly samples

For analysis of dSarm expression, fly heads were collected from 1-day old flies at 5-days post rotenone treatment in the presence or absence of Olaparib and crushed in a mortar-pestle with TRIZOL. The cDNAs were prepared by iScript cDNA synthesis kit from Bio-Rad and the mRNA expressions were measured by drosophila specific primers by ΔΔCT method as discussed in above and analyzed by CFX manager (Bio-Rad). All results were collected from at least three independent samples.

### Statistical Analysis

All statistical analysis was performed using GraphPad Prism version 7.00 for Windows (GraphPad Software, La Jolla, CA, USA). The results are presented as mean ± SEM (standard error of mean) of at least three independent experiments performed in triplicate. Statistical significance was measured by one-way analysis of variance (ANOVA) with Bonferroni posttest. Asterisks indicate levels of significance (*p <0.05, **p <0.01 and ***p < 0.001).

## Acknowledgements

The authors would like to thank Rupak Datta, Abhik Saha and Shubhra Majumder for their insightful comments during the preparation of the manuscript and for providing several reagents. The work was funded by extramural funding from Dept. of Biotechnology, India (#BT/PR10983/BRB/10/1282/2014) to PM. AS is a recipient of UGC-NET fellowship [F.16-6(Dec.2016)/2017(NET)] and SD is a recipient of CSIR-NET fellowship [08/155(0085)-2020-EMR-I] from Govt. of India.

## Author contribution

PM conceived the work and AS, MS, SD and PD performed the experiments. AS and PM wrote the manuscript and prepared the figures and PM revised the entire manuscript.

## Supplementary figure legends

**Fig. S1.**
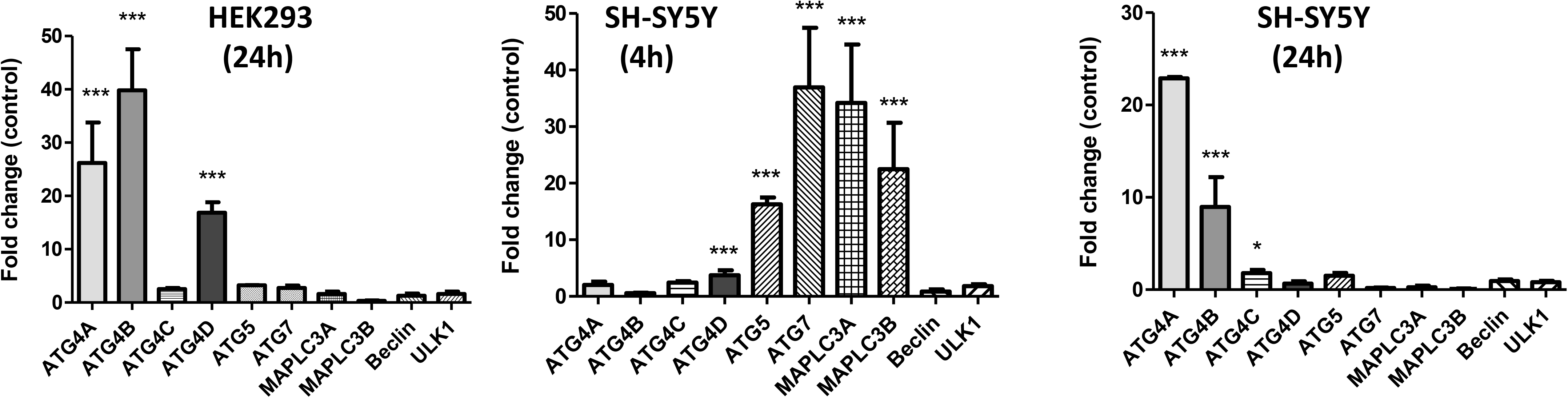
(A-C). qRT-PCR analysis of the fold change in gene expression of selected autophagy genes at 4 or 24 h post rotenone treatment in HEK293 (A) and SH-SY5Y (B-C) cells with respect to DMSO treated controls. Gapdh was used as an internal control for the analysis.

**Fig. S2.**
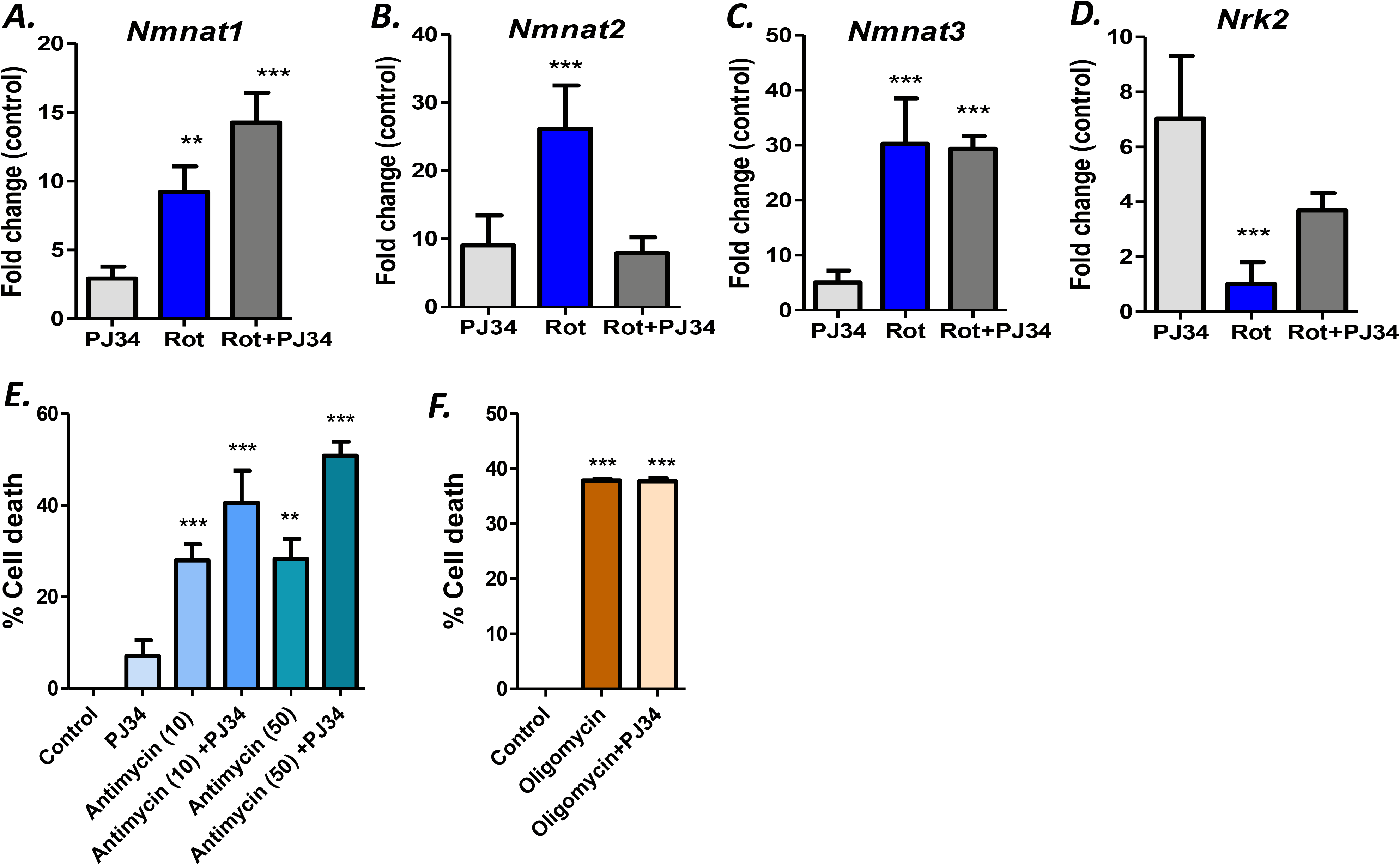
(A-D). qRT-PCR analysis of the fold change in gene expression of Nmnat1 (A), Nmnat2 (B), Nmnat3 (C) and Nrk1 (D) at 24 h post rotenone treatment in HEK293 with respect to DMSO treated controls. Gapdh was used as an internal control for the analysis **(E-F)** Cell viability assay in HEK293 cells pre-incubated with 25 µM PJ-34 for 1 h followed by treatment with the mitochondrial complex III inhibitor Antimycin A (10 µM and 50 µM) (E) and the mitochondria complex V inhibitor Oligomycin (500nM) (F). The percentage of cell death was determined by the OD_540_ values in an MTT assay.

**Fig. S3.**
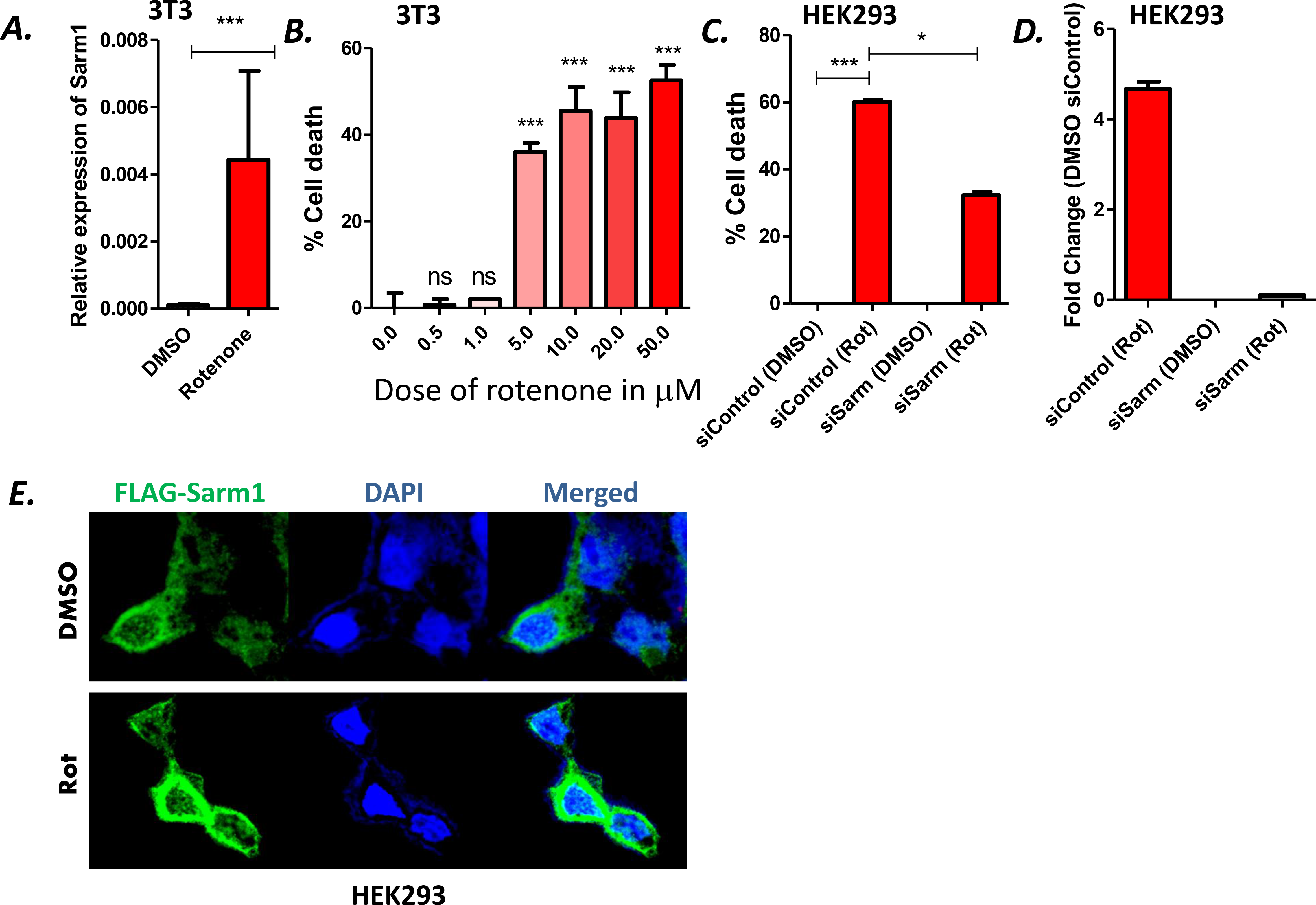
(A). qRT-PCR analysis of Sarm1 gene expression at 24 h post rotenone treatment in 3T3 cells. Gapdh was used as an internal control **(B).** MTT assay of cells treated with indicated doses of rotenone at 24 h post treatment in 3T3 cells **(C).** Cell viability assay in Sarm1 knock-down HEK293 cells followed by rotenone treatment for 24 h and the results were compared with control siRNA samples. **(D)** qRT-PCR analysis of the fold change in gene expression of Sarm1 at 24 h post rotenone treatment in HEK293 with respect to DMSO treated controls in both siControl and siSarm1 samples. Gapdh was used as an internal control for the analysis. Data are representative of two independent experiments with three replicates per sample **(E).** Immunofluorescence analysis of HEK293 cells transfected with FLAG-tagged full length Sarm1 (Green) for 48 h followed by rotenone (500 nM) treatment for 24 h. Results were compared with DMSO treated but transfected cells. Nuclear staining is represented with DAPI (blue).

**Supplementary Table1:**
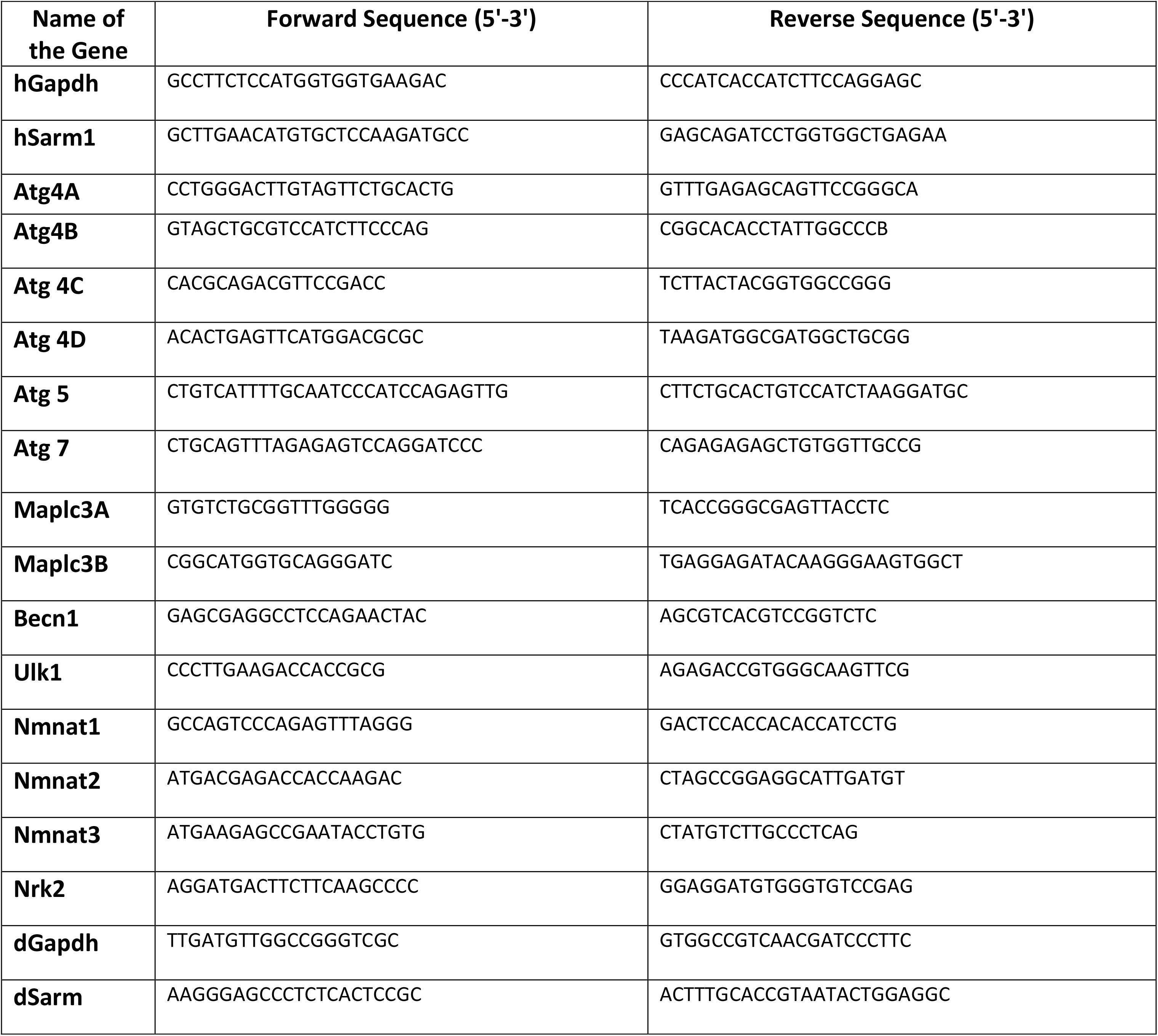

